# Quantitative genetic architecture of adaptive phenology traits in the deciduous tree, *Populus trichocarpa* (Torr. & Gray)

**DOI:** 10.1101/2020.06.12.148445

**Authors:** Thomas J Richards, Almir Karacic, Rami-Petteri Apuli, Martin Weih, Pär K. Ingvarsson, Ann Christin Rönnberg-Wästljung

## Abstract

In a warming climate, the ability to accurately predict and track shifting environmental conditions will be fundamental for plant survival. Environmental cues define the transitions between growth and dormancy as plants synchronise development with favourable environmental conditions, however these cues are predicted to change under future climate projections which may have profound impacts on tree survival and growth. Here, we use a quantitative genetic approach to estimate the genetic basis of spring and autumn phenology in *Populus trichocarpa* to determine this species capacity for climate adaptation. We measured bud burst, leaf coloration, and leaf senescence traits across two years (2017-2018) and combine these observations with measures of lifetime growth to determine how genetic correlations between phenology and growth may facilitate or constrain adaptation. Timing of transitions differed between years, although we found strong cross year genetic correlations in all traits, suggesting that genotypes respond in consistent ways to seasonal cues. Spring and autumn phenology were correlated with lifetime growth, where genotypes that burst leaves early and shed them late had the highest lifetime growth. We also identified substantial heritable variation in the timing of all phenological transitions (h^2^ = 0.5-0.8) and in lifetime growth (h^2^ = 0.8). The combination of abundant additive variation and favourable genetic correlations in phenology traits suggests that cultivated varieties of *P. trichocarpa* have the capability to create populations which may adapt their phenology to climatic changes without negative impacts on growth.

## INTRODUCTION

Perennial plants transition between periods of growth and dormancy in response to seasonal changes. Patterns of growth cessation and dormancy minimise the risk of tissue damage from freezing over winter and are primarily induced by a change in photoperiod (Fracheboud *et al.* 2009; Basler and KÖrner 2012). Once dormant, many plants require a period of chilling before active growth is resumed in response to warming spring temperatures (Horvath *et al.* 2003). These transitions define the trade-off between growth and potential damage and are therefore expected to be subject to natural selection. Widespread species are exposed to a range of temperature and photoperiod length across their distribution, leading many species to show local adaptation and heritable variation in the timing of these phenology transitions associated with local conditions (Howe *et al.* 2003; Luquez *et al.* 2008; Cong *et al.* 2016). The adaptive significance of cyclic phenology is especially important in woody perennials such as trees, as they are long lived and therefore exposed to seasonal changes over multiple years.

While both spring and autumn phenology determine growing season and are therefore potentially adaptive, transitions are determined by different cues and therefore may respond to selection in different ways (Singh *et al.* 2017). Dormancy is broken by changes in temperature at the beginning of spring in many plants (McKown *et al.* 2018), and therefore bud burst is defined by cues which have significant variation between local environments and successive years (Basler and KÖrner 2012; Brelsford *et al.* 2019). Growth cessation is generally induced by changes in photoperiod (Fracheboud *et al.* 2009; Lagercrantz 2009; Michelson *et al.* 2017) or associated changes in the light spectrum (Leuchner *et al.* 2007; Brelsford *et al.* 2019), and therefore varies predictably with latitude and consistently between growing seasons. These different characteristics between the triggers of growth and dormancy mean that the adaptation toward the optimal conditions and timing for spring and autumn phenology requires different biological responses. Consequently, effects of natural selection on genetic variation in these traits is likely to differ, potentially driving differences in the genetic architecture underlying each trait and influencing the scope of adaptive shifts in phenology traits under novel climatic conditions.

Phenotypic variation is well characterised for many physiological and phenological traits in deciduous trees (Costa e Silva *et al.* 2004; Fabbrini *et al.* 2012), foremost of which are the Poplars, due to their adaptability, widespread distribution and commercial value. A number of studies have investigated the genetic basis of poplar phenotypes and have identified many genomic variants associated with phenology and growth (Ingvarsson *et al.* 2008; Luquez *et al.* 2008; Fabbrini *et al.* 2012; Triozzi *et al.* 2018). However, adaptability in complex traits largely depends on the pleiotropic effects of this genetic variation which can manifest as genetic covariance between phenology and growth-related traits (Porth *et al.* 2014; McKown *et al.* 2018). This information is fundamental to understanding the adaptability of poplar species, as the degree of genetic covariance between phenology traits and growth ultimately determines the magnitude of response to selection on individual traits (Porth *et al.* 2015). A life history strategy that minimises the risk of cold damage through late bud burst and early bud set may reduce the overall growth and therefore constrain growth and establishment. Similarly, a strategy of early bud burst and late bud set results in a long growing season but increases the risk of frost related injury. As such, the capacity of trees to adapt to climatic variation is dependent on the genetic variance and covariance of phenology traits and growth, which determines the current range limit (Chuine 2010) and potential to expand into novel habitats (Zanewich *et al.* 2018).

*Populus trichocarpa* (black cottonwood) is one of several large, deciduous, tree species within the genus *Populus. Populus* species are generally fast-growing and as such have great ecological significance as pioneer species (Cronk 2005), as well as an increasing value as a species for short rotation forestry (Weih 2004) and bioenergy production (Sannigrahi *et al.* 2010; Porth and El-Kassaby 2015). As a result, there has been extensive development of genetic resources to investigate and capitalise on the genetic basis of adaptation and growth traits in these species (Cronk 2005; Jansson and Douglas 2007). Extensive research has identified substantial genetic variation across several adaptive traits, and while these estimates derive from a range of experimental designs and study populations, moderate to high heritability has been generally reported for growth (Bradshaw and Stettler 1995; Yu *et al.* 2001; Porth *et al.* 2015), leaf phenology (Howe *et al.* 2000; McKown *et al.* 2014a; Porth *et al.* 2015) and cold tolerance (Howe *et al.* 2003) traits. A distribution over a wide range of geography and climate has driven patterns of local adaptation (Evans *et al.* 2014; McKown *et al.* 2014a), however extensive long distance pollen transfer and large population sizes have resulted in very low population differentiation across its natural range (Slavov and Zhelev 2010). Despite this, long running breeding programs have produced few varieties that are well suited to the photoperiod and climate typical of higher latitudes (Karacic *et al.* 2003) which may limit establishment of populations outside of the natural climatic range.

Here, we analyse phenotypic variation in a clonally replicated population of *P. trichocarpa* genotypes grown in central Sweden. We survey the genetic basis of spring and autumn phenology and determine the genetic covariance of these traits with fitness in the form of lifetime growth. These relationships determine the genetic constraints on adaptive evolution in this species and as such may constrain range expansion into hostile environmental conditions. Specifically, we 1) aim to characterise heritable genetic variation for phenology traits in this population, and 2) test whether genetic correlations that exist between phenology traits may facilitate or constrain adaptation to novel climatic conditions.

## MATERIALS & METHODS

### POPULATION CHARACTERISTICS AND PHENOTYPIC MEASUREMENTS

Here we present a quantitative genetics analysis of phenology measurements taken from 564 mature *P. trichocarpa* trees in a plantation at Krusenberg (59°44’44.2”N 17°40’31.5”E) in central Sweden. Material in this plantation was originally generated from 9 female and 10 male trees collected over a latitudinal range from 44-60° in North America (Supplementary Table 1), which were randomly crossed to produce 34 families. From these families, a total of 120 half sib, full sib, and unrelated trees (onward referred to as genotypes) were clonally replicated with between 1 and 20 (median = 6) individual trees per genotype represented in the study population. The plantation was established in 2003 on a flat, homogeneous area of agricultural field approximately 275m × 40m. Individuals were planted at 3.5 m quadratic spacing in a randomized design. The experiment was systematically thinned in March 2013 leaving 564 trees in an approximately 3,5×7 m diamond spacing.

**Table 1:**
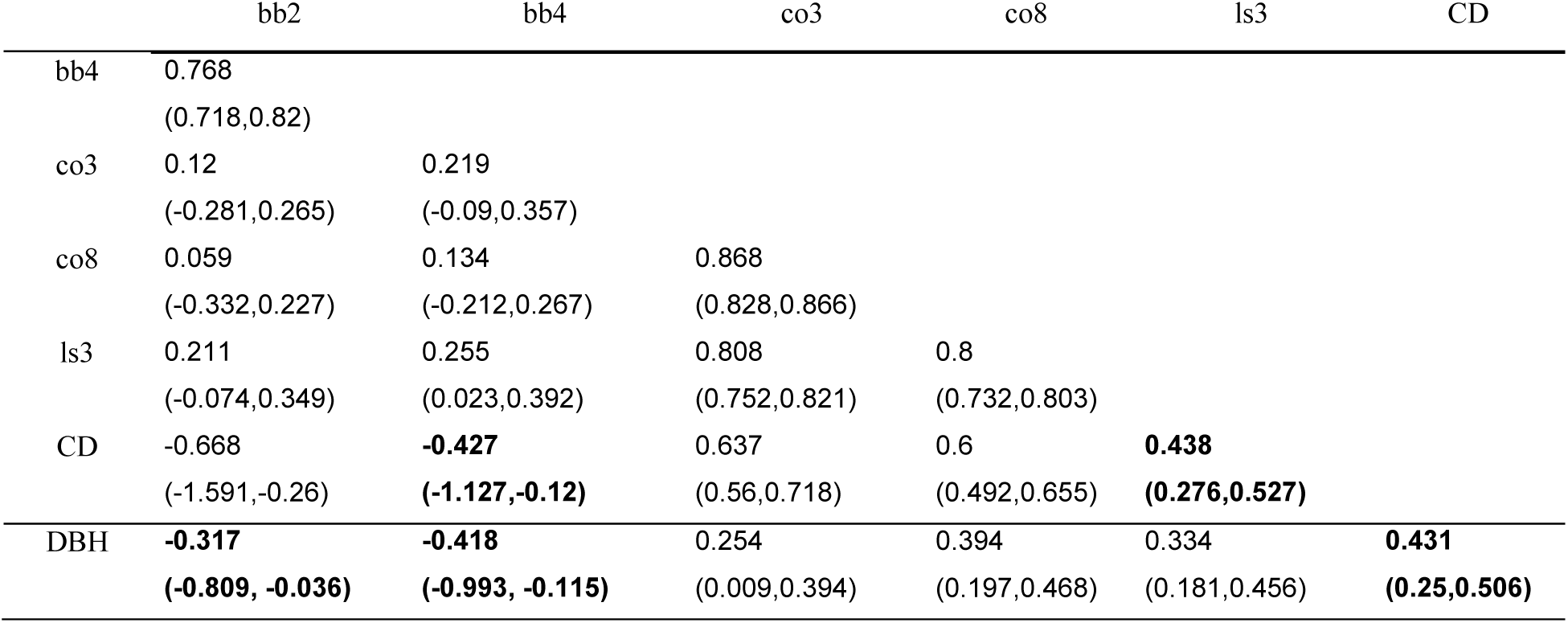
Genetic correlation matrix of phenology and growth traits in *P. trichocarpa.* Correlation estimates were drawn from a 7 trait, multivariate ‘animal’ model using the MCMCGLMM procedure in R. Values in brackets represent the lower and upper 95% credible intervals, values in bold do not overlap zero. Traits include beginning and end of budburst (bb2, bb4), beginning and end of leaf colour transition from green to yellow (co3, co8), leaf shed (ls3), canopy duration (CD), and lifetime growth (DBH).

To asses variation in phenology we included the traits that best define the major milestones of phenology during the annual growth cycle. For spring development, bud burst (bb) was scored on a scale from 1-5, with stage bb2 representing initial shoot emergence, bb3 leaf primordia exposed, bb4 leaves half shed with bud scales dropped and bb5 leaves completely shed. Autumn phenology was scored on a scale of 1-8 based on a continuous range of crown colouring from col1= 100% green, to col8 100% yellow. Leaf senescence (ls) was measured on a 3-point scale where ls1 = full foliage, ls2 = half leaves remaining and stage ls3 full defoliated. We estimated growing season length by the duration of photosynthetically active leaf canopy. Canopy duration (CD) was defined as the period between the beginning of budburst (bb2) and beginning of leaf yellowing (co3). Phenology measurements were taken every 2-5 days during the spring and autumn seasons in 2017 and 2018. Lifetime growth was determined by measuring diameter at breast height (DBH) in 2017

While late season growth cessation is best described by bud set, this trait is difficult to accurately measure in mature trees. Due to the difficulty of determining the exact transitions between beginning of growth, cessation of growth and bud development in grown trees, we split these seasonal transitions into 5 biologically relevant proxy stages which describe the start and end of seasonal transition; bb2, the first bud emergence, (bb5) full leaf emergence, (col2) first stage of yellowing, (col5) complete yellowing, (ls3) day of full leaf shed. These phenology traits are combined with growth of the tree at 14 years of age as measured by cross calliper measurement (DBH) in 2017.

### DATA IMPUTATION

Screening was conducted at intervals of 2-5 days meaning that individual trees occasionally passed through developmental stages between screenings. To account for these missing estimates of the day of transition we estimated the transition days for each developmental stage using local regression models (LOESS) fit to each individual tree. Models were fit through the data point of the first day an individual tree was observed at a stage transition and day estimates were calculated for any stage transitions for which there was no direct observation (Figure S2). This method estimates a non-linear developmental curve which is not constrained to fit any a-priori mathematic distribution for each individual tree and allows estimation of the day in which trees passed each developmental stage and inclusion of individuals which were not observed at important developmental transitions in the analysis. As extrapolation beyond the range of the observed data is unreliable using this method, we only retained estimates that were bounded by observations on either side. We validated this method by comparing estimated values with direct observations to ensure that estimated transitions accurately represented the observed data (Pearsons R = 0.98-1).

## STATISTICAL ANALYSIS

### Heritability and cross year genetic correlations

We estimated genetic parameters in this population using Bayesian Mixed model approach implemented in the MCMCglmm R package (Hadfield 2010). This approach uses pedigree information to construct a relatedness (A) matrix which allows for a proper structuring of the random genetic effects to account for the complex combination of unrelated individuals, with half and full sibs included in the plantation. To estimate additive genetic heritability (*h*^*2*^) for each genotype, and to determine the additive genetic co-variance between measurement years, we implemented the following bivariate models, where measures of each transition trait (e.g., budburst) taken in subsequent years 2017 *(Y1)* and 2018 *(Y2)* formed a bivariate response variable, *µ* is a fixed intercept, g is the random additive genetic component drawn from pedigree information and *ε*_*ijk*_ residual variance. 

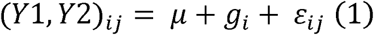

Following the approach of Kennedy and Schaeffer (1989), we treat genetically identical individual trees (clones) from the same genotype as repeated measures, and estimate the genetic parameters based on the definition of the relatedness matrix (A) among genotypes rather than cloned individuals. Variation between individual trees within genotype was partitioned into the residual variance. Each trait was modelled separately. Models were run for 1000000 iterations, and we implemented Markov chain Monte-Carlo sampling with a thinning interval of 1000 to ensure very low autocorrelation between samples (> 0.003), and burn-in period of 1000 iterations. This approach yielded posterior probability distributions with sample sizes for each trait pair near 1000, from which we derived parameter estimates with 95% credible intervals. In all cases we specified uninformative priors (variance = 1, degree of belief = 0.002), although we also ran models with a range of prior specifications to ensure the final parameter estimates were not affected by our chosen priors. We extracted estimates for genetic effects from the posterior distribution of model (1) and calculated narrow sense heritability by extracting the additive genetic (Va) proportion of total trait variance (Va + Ve). Note that we estimate heritability based on each genotype, with variance among individual trees within genotype partitioned in the residual term.: 

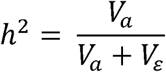

Separate heritability estimates were calculated for each phenology trait in two consecutive seasons (2017 and 2018). We do not directly account for dominance effects in this model due to limited size and depth in the pedigree, and as such our estimates of heritability may be overestimated as some portion of any potential dominance variance may contribute to the estimates of *V*_*a*_ (Walsh and Lynch 2018). Cross-year genetic correlations were calculated by extracting the 2×2 genetic variance-(co)variance matrix from model (1), and we calculated genetic correlations as the genetic covariance between traits divided by the square root of the product of the variances.

## GENETIC CORRELATIONS OF SPRING AND AUTUMN PHENOLOGY

We calculated the additive genetic correlations between phenology transitions and lifetime growth to determine the genetic architecture relating phenology to fitness. We combined the phenology transitions of budburst (bb2), leaf expansion (bb5), start of leaf yellowing (col2), end of leaf yellowing, (col5) leaf shed (ls3), canopy duration (CD) and lifetime growth (DBH) as a 7-trait multivariate response (*Y*_*ijk*_), and implemented the following model: 

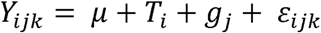

Where *µ* is a fixed intercept, *T*_*i*_ is the fixed effect of year, *g*_*j*_ is the random term describing the additive genetic variance drawn from pedigree information and *ε*_*ijk*_ residual variance. We implemented this model using a Bayesian ‘animal’ model approach in the MCMCGLMM package in R (Hadfield 2010). We implemented Markov Chain Monte Carlo sampling to estimate posterior probability distributions for the genetic parameters, and increased iterations until sample sizes for each estimation approached 1000. Our final models included 500,000 iterations, with a thinning interval of 500 and a burn-in of 10,000 iterations. Priors were constructed using measured phenotypic variances with a low degree of belief, and were checked against uninformative priors with low and high degrees of belief to ensure that prior specification had little effect on parameter estimates.

We extracted the 7×7 genetic variance-(co)variance matrix from this model, and calculated genetic correlations as the genetic covariance between traits divided by the square root of the product of the variances. We extract the genetic correlation between the 6 phenology traits and fitness (measured as DBH17, or lifetime growth) to determine the genetic relationship between phenology and growth. This correlation between phenology and growth ultimately determines whether adaptive shifts in phenology in response to climate change will be associated with increased growth, or be constrained by associated negative effects on tree growth.

## RESULTS

### Phenotypic distribution, summary statistics

All phenology transitions occurred earlier in 2018 than 2017 (Figure 1). Differences in the mean date for each stage were between 3 days (leaf shed, bud burst) and almost 10 days (full yellow) earlier in 2018. Spring transitions occurred faster in 2018; bb2 occurred over a period of 39 days in 2017 and 17 days in 2018 (difference of 22 days), while leaf out (bb4) took 17 days in 2017 and 8 in 2018 (difference of 9 days). These differences corresponded with temperature and rainfall variation between observation years, where temperature increase was more gradual during spring in 2017, summer was warmer and wetter in 2018, and autumn was drier in 2018. The duration of spring transitions was also shorter in 2018 indicating the abrupt increase in spring temperature during that year sped up leaf unfurling. (Figure S1).

**Figure 1:**
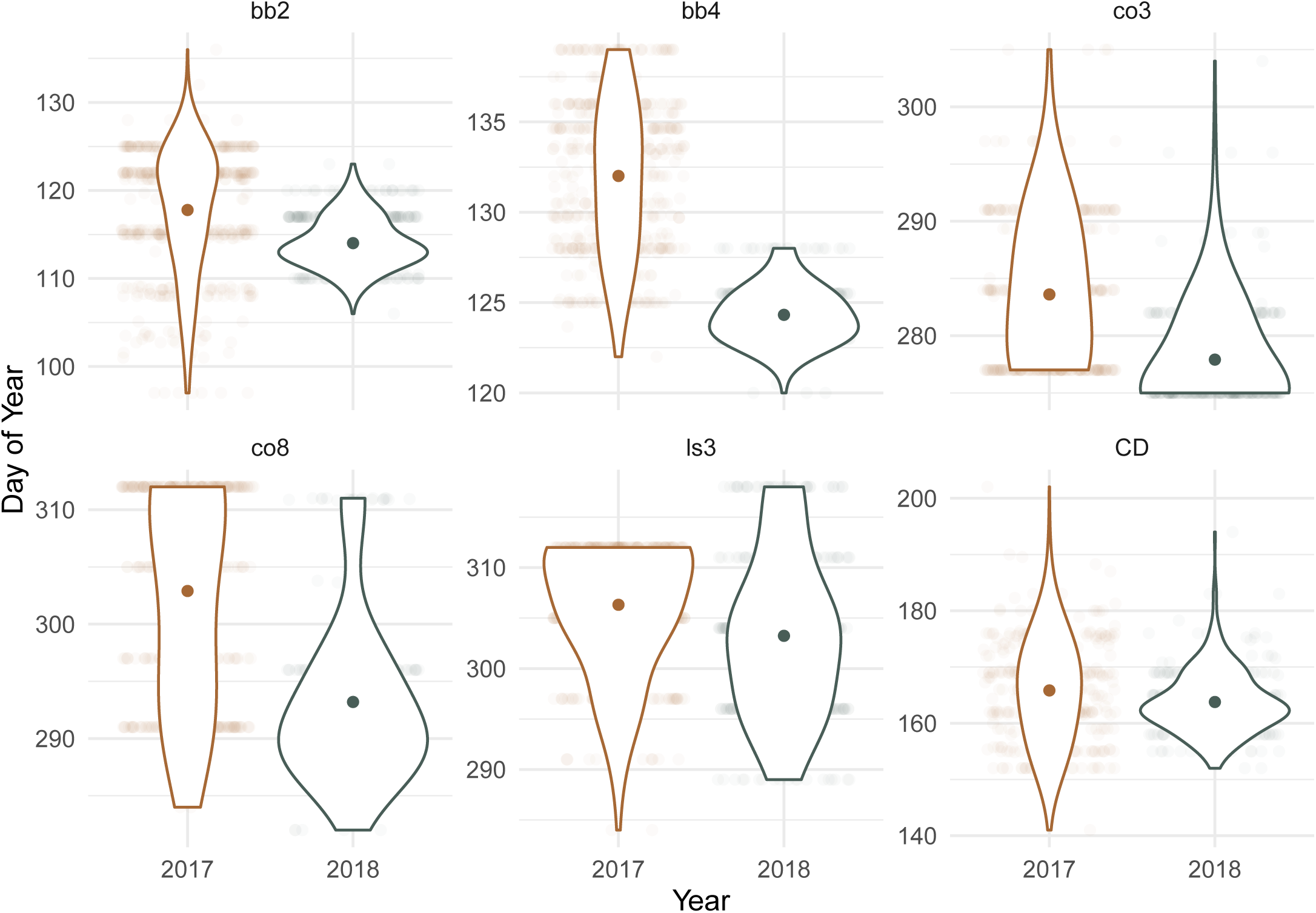
Phenotypic trait summary in 2017 and 2018. Summaries shown are mean and density overlayed upon original data points. All phenology transitions occurred earlier in 2018 leading to a slightly longer mean growing season. Truncated distributions are indicative of an abrupt timepoint where all individuals reached maximum scores in each trait. Traits include beginning (bb2) and end (bb4) of budburst, beginning (col3) and end (col8) of leaf colour transition from green to yellow, leaf shed (ls3), canopy duration (CD, and lifetime growth (dbh).

### Trait heritability

All phenology traits had moderate to high heritability with median estimates in the range of 0.41-0.76, identifying significant genetic variation in phenology traits between the genotypes in this growth trial (Figure 2). Heritability was highest in budburst (2017), leaf shed (2017, 2018) and lifetime growth traits, and lowest for leaf out and colouration transitions. Estimates for 2018 are reduced across all traits except leaf shed, although there is considerable overlap in posterior distributions for estimates in all traits except budburst.

**Figure 2:**
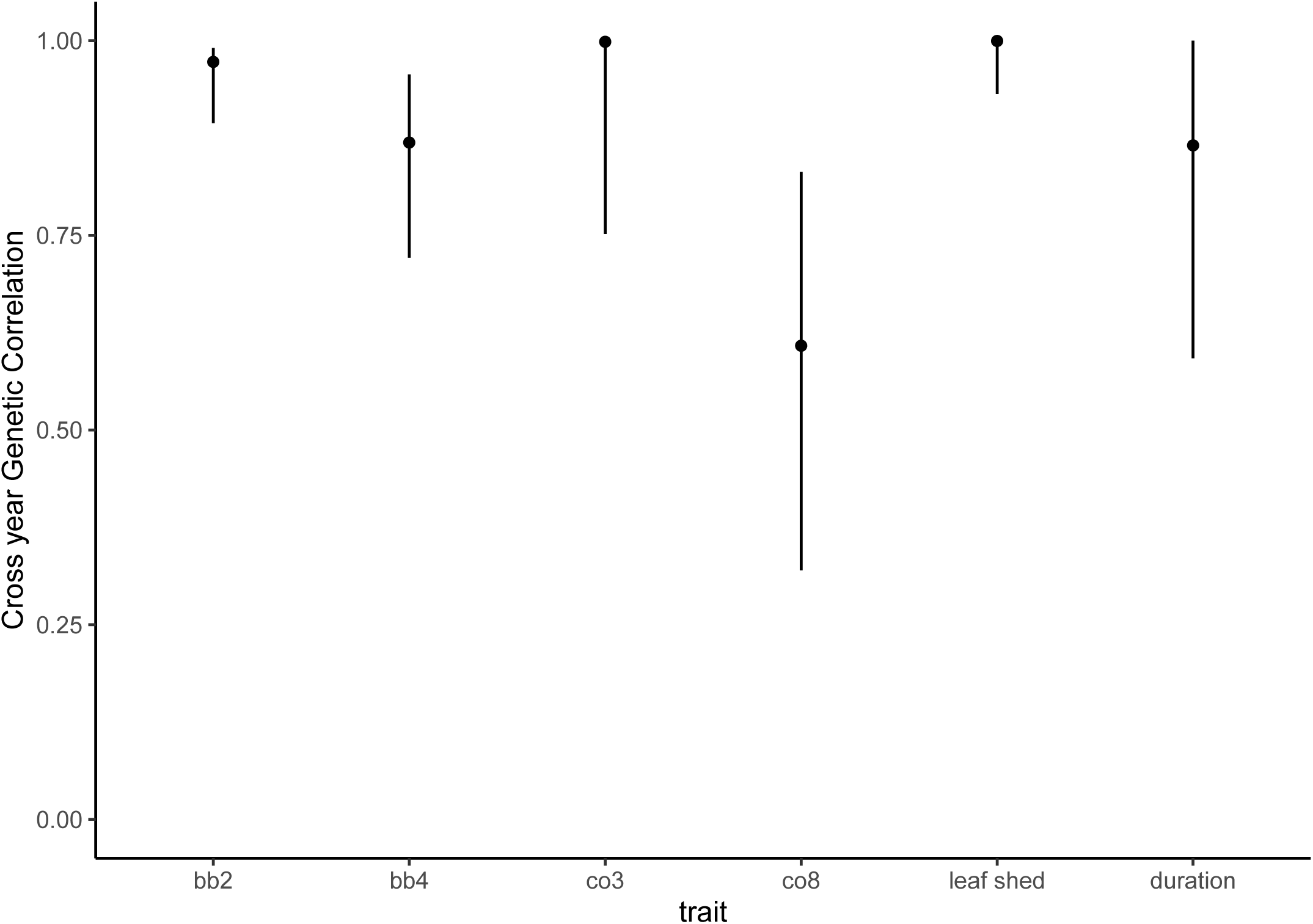
Narrow sense heritability estimates for phenology traits at Krusenberg field experiment in 2017 and 2018. Means and 95% credible intervals derived from univariate ‘animal’ models run in R (MCMCglmm)

**Figure 3:**
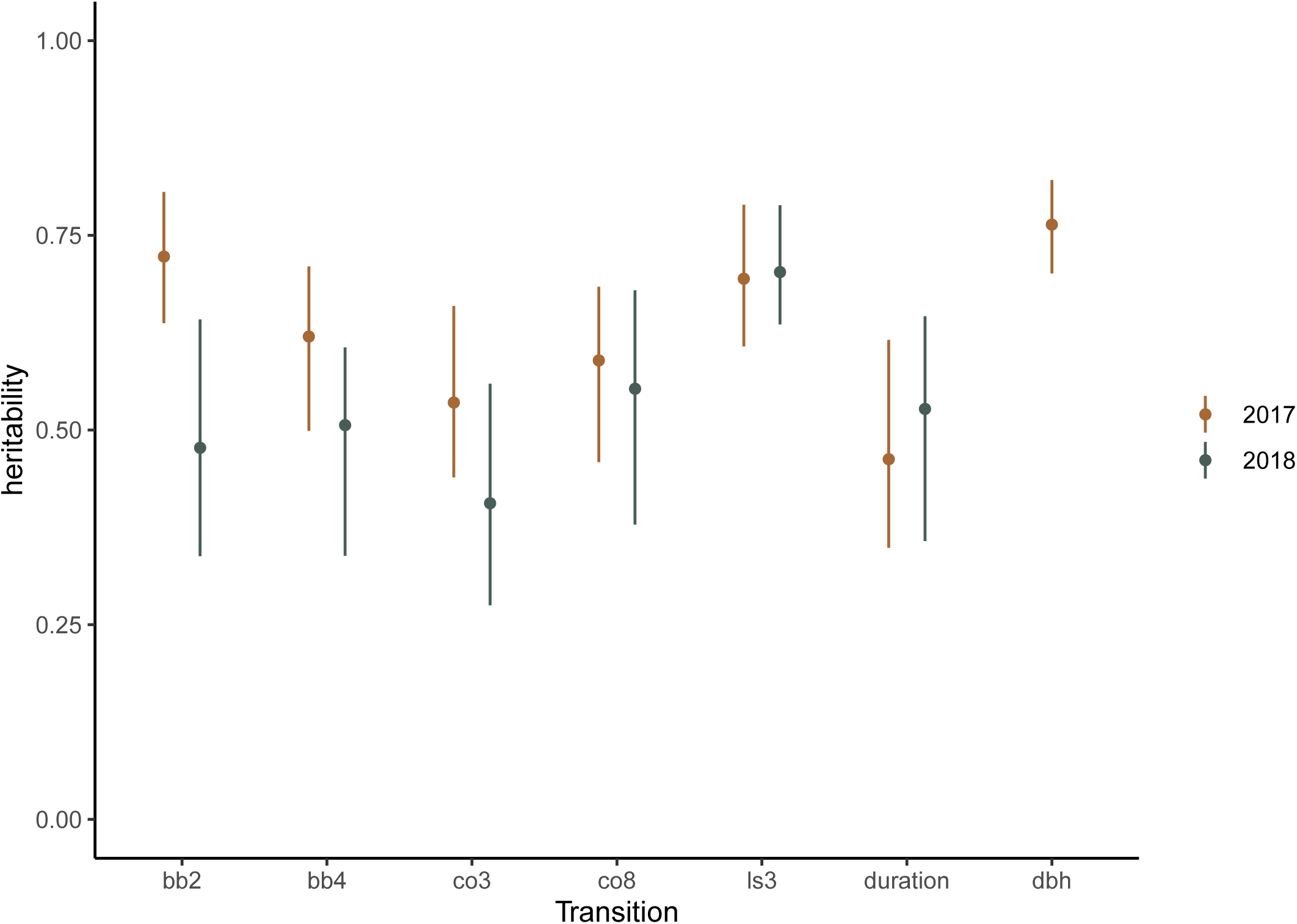
Cross year genetic correlations. Mean and 95% credible intervals for estimates of cross year genetic correlations. Traits include beginning and end of budburst (bb2, bb4), beginning and end of leaf colour transition from green to yellow (co3, co8), leaf shed (ls3), canopy duration (CD), Correlations between years is consistently high, and intervals overlap 1 for bb2, co3 and leaf shed traits.

Phenology traits had a strong genetic basis, with spring heritability in the range of 0.48 - 0.72 (bb2) and 0.51 - 0.62 (bb4) respectively. Autumn phenology was also highly heritable, with highest and most consistent estimates for leaf shed (ls3) (*h*^*2*^ = 0.60 – 0.64). Lifetime growth was the highest and most precise of the heritability estimates (*h*^*2*^*=*0.76), which reflects a substantial genetic basis for differences in lifetime growth between genotypes.

### Cross year genetic correlations

Despite the considerable interannual shifts in phenotypic means described above, cross year genetic correlations were high. Estimates for budburst (bb2), onset of colour change (col3), and leaf shed (ls3) all overlap 1, showing complete correlation between years. The contribution of genetic variation to leaf out (bb5) and canopy duration also very consistent with correlations above R = 0.8. The transition to full yellow leaves cross-year correlation was R = 0.6, which is high given that this trait also had the largest mean difference between years at 9.7 days.

### Additive genetic correlations between traits

Spring traits of budburst (bb2) and leaf out (bb5) displayed highly positive genetic correlations (R = 0.77), but showed little to no link to autumn phenology characteristics (Table 1). Similarly, autumn traits were strongly correlated indicating strong alignment in the genetic basis of traits within either spring or autumn seasons but not between. Leaf colouration was highly correlated with leaf senescence (R = 0.80-0.87) suggesting a developmental cascade brought on by pleiotropic effects of genes involved in sensing and responding to the end of favourable growing conditions. Spring phenology was negatively correlated with both canopy duration and lifetime growth, where clones bursting leaves early in the season hold their leaf canopy longer and grow more. Leaf colouring was positively correlated with growth reflecting that clones with late onset of leaf colouration and senescence conferred higher lifetime growth (Table 1).

## DISCUSSION

The ability of plants to survive the novel environmental conditions brought on by climate change is dependent on populations harbouring sufficient genetic variation to adapt to changes in seasonality. Here, we show that timing of phenology in an experimental population of *P. trichocarpa* has a heritable genetic basis, but also that these traits can vary between years in response to climatic variation. With heritable genetic variation underlying both spring and autumn phenology, and an absence of genetic constraint between these transitions, our results suggest that these traits have potential to evolve independently in order to track future climate changes. As projections suggest climate change may provide conditions for a longer growing season, the absence of constraint may allow trees to have a longer canopy duration, which will in turn lead to higher growth. Together these findings provide insight into the genetic architecture of phenology traits, and identify the capacity of *P. trichocarpa* to adapt its phenology to novel environmental contexts as they colonise areas outside their original range, or as environmental conditions shift via global climate change.

There is a considerable contribution of genetics to all phenology traits surveyed in this field experiment and similar to other studies (eg. Olson *et al.* 2013) we find abundant genetic variation for phenology traits determining the length of the growing season. There was also a strong genetic correlation within spring and within autumn phenology traits (0.77 - 0.87) suggesting pleiotropic effects of the genes underlying the cascade of leaf development and between traits influencing leaf coloration and leaf senescence, however genetic correlations across spring and autumn phenology were low to non-existent, reinforcing previous studies that show the genes (McKown *et al.* 2014b) and environmental cues (Singh *et al.* 2017) underlying spring and autumn phenology are different. Both spring phenology traits had negative correlations with lifetime growth suggesting that earlier budburst is beneficial for lifetime growth, and there was a stronger link between leaf unfurling and growth, suggesting clones with predictable late leaf unfurling was detrimental for lifetime growth.

Growing season in deciduous forest trees is defined by the leaf phenology traits, and we show with this experiment that canopy duration has a negative genetic correlation with spring phenology, and a positive correlation with autumn phenology. Perhaps unsurprisingly, we identify a genetic association where early bud burst and late leaf drop leads to longer growing season and greater lifetime growth. These patterns have been shown in other studies of this species (eg. McKown *et al.* 2014a), reflecting the general biology of adaptation in deciduous plants, where the timing of spring phenology resolves the tension between two factors: damage avoidance and competition. Early budburst increases the probability of leaf exposure to frost conditions, however it may confer a competitive advantage by earlier exposing leaves to (direct) sunlight. Late leaf flush delivers trees into an environment where much of the available light is already absorbed by early flushing neighbours, and puts the tree at a disadvantage for light capture that may last for the full growing season. This disadvantage likely accumulates across years as the late flushing individual falls further behind its faster growing neighbours. Late flush also reduces exposure to optimal spring growing conditions, further compounding the disadvantage of spring shading which may significantly reduce growth (Yu *et al.* 2001).

A number of studies have found moderate heritability for phenology traits in this, and other *Populus* species for estimates of bud flush however, it is important to note that the majority of these studies have been conducted on juvenile trees (eg. McKown *et al.* 2014a; Pliura *et al.* 2014). Estimates of growth heritability presented here is generally higher than previously reported values for *P. trichocarpa* (Marron *et al.* 2010; McKown *et al.* 2014a), however this may be due to estimations being derived from established adult trees which will reduce the impact of random events which can lead to highly variable growth in the first few growing seasons. It is not uncommon to find higher heritability for fitness related traits in older individuals in animals and humans (eg. Christensen *et al.* 2003; Wilson *et al.* 2005) however this is more likely to be due to a decline in environmental or residual variance, rather than an increase in additive genetic variance. Other considerations such as the pedigree structure and source material in this population may also contribute to the higher heritability estimates than other studies, and also contribute to the large credible intervals around these estimates. We emphasise that due to these factors, heritability estimates should be interpreted in the context of the present population and experimental design.

While we find evidence for a heritable basis to spring and autumn phenology, there is only a weak relationship between leaf unfurling and senescence. This finding adds to existing evidence that different genetic architectures are controlling phenological responses of spring and autumn environmental cues in *P. trichocarpa*. This is consistent with the observation of conserved patterns of environmental responses across the plant kingdom where bud burst is largely regulated by local environment not local adaptation (MacKenzie *et al.* 2018) and leaf out highly dependent on temperature (Polgar and Primack 2011). Sensitivity to local growing conditions such as temperature and altitude (a strong determinant of temperature) can be stronger than population level patterns of local adaptation, and sensitivity can be similar between populations of the same species (Vitasse *et al.* 2009), suggesting that the timing of spring growth is mediated by plants anticipating favourable local conditions which pose reduced risk of damage before investing resources in the production of leaf tissue. The strong cross-year genetic correlations and high heritability in these traits suggests that there is substantial variation in the timing of seasonal transitions across years, but that clones respond in the same way over consecutive years.

Variability and uncertainty in relation to the future timing of seasonal transitions and the potential for extreme or unseasonal weather events such as out of season frost pose a significant risk of damage to deciduous trees. Our findings here suggest that adaptive shifts in phenology may be possible in *P. trichocarpa*, however the genetic links between phenology and growth suggest that shifts in the timing of spring and autumn transitions have the potential to affect growth if shifts lead to a shortening of the growth period. Previous work has shown that autumn phenology is primarily initiated as trees respond to predictable changes in photoperiod, but data presented here suggests that large interannual shifts in autumn timing are possible, which suggest that leaf colour and senescence traits may also be responding to multiple environmental cues either during the summer growing season, or during the onset of autumn (Rohde *et al.* 2011b).

Multiple studies have shown that temperature is not a trigger for senescence (Bhalerao *et al.* 2003; Keskitalo *et al.* 2005; Luquez *et al.* 2008; Fracheboud *et al.* 2009), however speed of senescence is temperature dependent once initiated (Fracheboud *et al.* 2009) and evidence from poplar hybrids suggests that temperature may contribute to the timing of growth cessation and bud set (Rohde *et al.* 2011a). In this study we were unable to accurately measure bud set across the experiment, however it is clear that growth cessation occurs some time before bud set and before leaves change colour (Rohde *et al.* 2011b) which suggests that the annual variation in colour timing is a response to differences in autumn temperature between years that is occurring after the point where trees have initiated senescence. The risks of autumn frost damage are elevated with late bud set, which can have significant negative effects on viability and winter survival in juvenile trees (Howe *et al.* 2003), so it is likely that late season phenology is strongly associated with climatic adaptation. In this case, photoperiod is a more reliable predictor of the onset of frost than temperature, which fluctuates between seasons so it is likely that growth cessation and bud set occurs before the prospect of frosts and trees are prepared for winter long before the leaf colour changes (Basler and KÖrner 2012).

It is important to note that a substantial shift in autumn phenology occurred between the two years of observation. In 2018, there was a faster increase in temperature and more sunny days during early spring, which is reflected in the faster spring development and earlier leaf out in that year. Autumn was also drier, and leaf colouration earlier in 2018, which is consistent with the presence of drought induced bud set, so it is likely that although light characteristics are the primary cue for growth cessation and senescence (Michelson *et al.* 2017; Triozzi *et al.* 2018), the timing and response to these factors is modified by other environmental cues such as temperature (Rohde *et al.* 2011a) or drought stress (Adams *et al.* 2015). Importantly, this finding suggests that although timing of phenology transitions have a strong genetic basis, trees respond to multiple cues and may respond to variable local climatic conditions as well as predictable cues such as day length. Adaptation may be facilitated by the capacity to track both constant cues (such as daylength) and variable cues such as temperature and rainfall, which may facilitate establishment of new populations outside of their natural range, and allow adaptation to shifting conditions within the native distribution.

## CAVEATS AND CONCLUSIONS

This study presents data from a collection of source genotypes sampled over approximately 15 degrees of latitude in north-western north America. This sampling design may inflate estimates of heritability by sampling across locally adapted populations, although this does not undermine the key findings that phenology transitions have a strong genetic basis. Also, we are unable to explore the contribution of dominance and epistatic interactions to our estimates of additive genetic variance due to limitations of the pedigree, however we note that these are common limitations of quantitative genetic analyses of experimental systems, and we acknowledge that some of the phenotypic variance attributed to additive genetic variation may be the result of dominance and epistatic interactions.

Overall, the substantial heritability found in phenology traits and the ability of trees to shift the timing of phenology transitions drastically between years suggests that *P. trichocarpa* populations have the ability to adjust their phenology in response to a changing climate. The wide and environmentally variable native range of *P. trichocarpa* is indicative of this species ability to persist across variable and unpredictable climates. We show here that a genetic variation underlies a large part of variation in phenology traits, but that trees also have the capacity for significant phenology shifts in response to year to year weather variation across the two years investigated here. This suggests this species has capacity to adapt its phenology to rapid climatic shifts, and that there is sufficient genetic variation underlying these traits that either natural or artificial selection could lead to evolutionary changes in adaptive phenology traits which may facilitate colonisation outside its natural range.

## Supporting information

Figure S2: Comparison of observed data (red) and Loess fit imputation (orange) for spring leaf development in four randomly selected individuals. Impu

Figure S1: Weather variation between the two years of survey. Spring temperatures increased faster, summer was warmer and wetter, and there were more

## ACKNOWLEDGEMENTS

This project was funded by the Swedish Research Council FORMAS, as part of the Climate-Adapted Poplar (CLAP) project.

## DATA ARCHIVING

Data will be uploaded to the Swedish Artportalen /Artdatabanken database, and any other archive suggested by Reviewers or editors

## FIGURE LEGENDS

**Figure S1**: Weather variation between the two years of survey. Spring temperatures increased faster, summer was warmer and wetter, and there were more sunny days during 2018. Historical range is indicated by black dots. Lines link monthly means for the study period.

**Figure S2:** Comparison of observed data (red) and Loess fit imputation (orange) for spring leaf development in four randomly selected individuals. Imputed data is fit through the first day individual trees were scored for each stage and show trajectories more in line with biological development. Intersection between dotted lines and fitted lines denotes values of bud break (BB2) and leaf unfurling (BB4).

**Table S1**. Crossing scheme showing origin (when known) of parents and number of clones from each cross. Some of the parents (in bold) and all offspring clones are included in the Krusenberg field experiment. OP indicates crosses that were conducted with Open pollination. Numbers within table indicate how many offspring were produced from each cross.

