## Supplemental Data 1 for "Quantitative genetic architecture of adaptive phenology traits in the deciduous tree, *Populus trichocarpa* (Torr. & Gray)"

| | | | Fathers | | | | | | | | | |
|---|---|---|---|---|---|---|---|---|---|---|---|---|
| | | Region | Alaska | | BC | Idaho | Oregon | Washington x Oregon | Washington | OP, Koster, Holland | | - |
| Mothers | | Lat (alt.) | 58°30' (15) | 60°00' | 49°15' | 48°25' | 45°35' | | 45°50' (75) | | | |
| Region | Lat (alt.) | Klon | **293** | **278** | **193** | 214 | 159 | 236 | **299** | 187 | 209 | 290 |
| Alaska | 58°30' | **216** | | | 5 | | 1 | | 5 | | | 5 |
| | 59°20' (500) | **291** | | | | | | 2 | | 1 | | |
| | 60°37' (400) | **265** | 3 | 3 | | 1 | | | 2 | 2 | | |
| BC | 49°00' | **230** | 7 | 4 | | 3 | | 2 | 6 | | | |
| | - | 96 | | | | | | | 1 | | | |
| Idaho x Oregon | - | 237 | 2 | 1 | | 3 | | | 3 | 1 | | 3 |
| | - | **240** | | 4 | | | | 1 | 21 | | 2 | |
| Oregon | 44°30' | 271 | 1 | 2 | | | | 1 | 4 | | | 1 |
| - | | 301 | | | | | | | | 2 | | 1 |
