## Supplementary figures and images for "Quantitative genetic architecture of adaptive phenology traits in the deciduous tree, *Populus trichocarpa* (Torr. & Gray)"

### Figure S1: Weather variation between the two years of survey. Spring temperatures increased faster, summer was warmer and wetter, and there were more

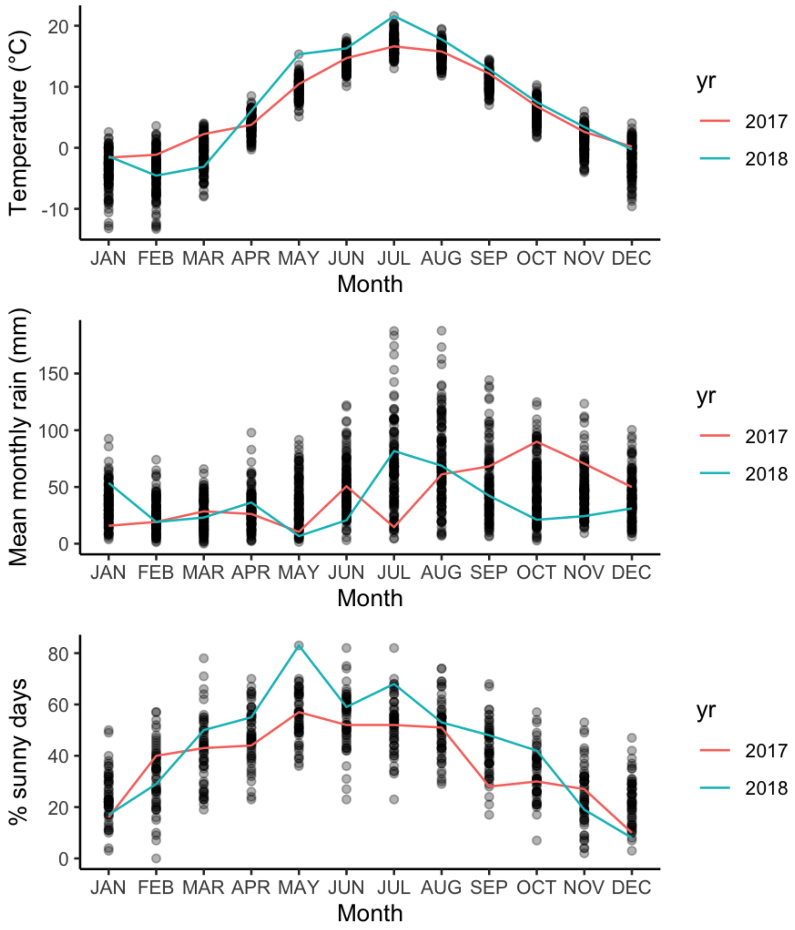

### Figure S2: Comparison of observed data (red) and Loess fit imputation (orange) for spring leaf development in four randomly selected individuals. Impu

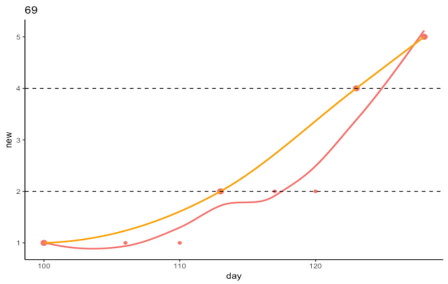

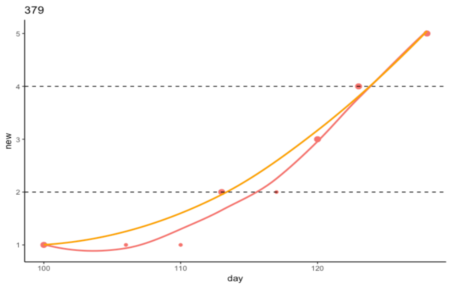

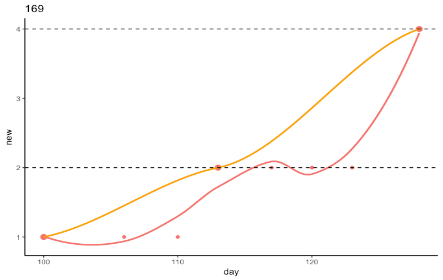

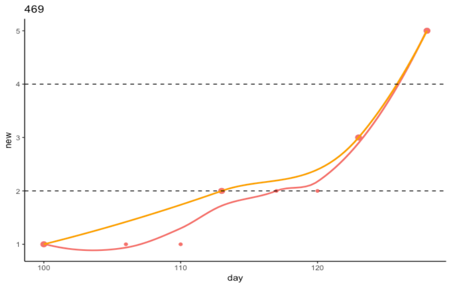
